# Masculinization of the X-chromosome in aphid soma and gonads

**DOI:** 10.1101/2021.08.13.453080

**Authors:** Julie Jaquiéry, Jean-Christophe Simon, Stéphanie Robin, Gautier Richard, Jean Peccoud, Hélène Boulain, Fabrice Legeai, Sylvie Tanguy, Nathalie Prunier-Leterme, Gaël Le Trionnaire

**Author notes:** **Cite as** Jaquiéry J, Simon J-C, Robin S, Richard G, Peccoud J, Boulain H, Legeai F, Tanguy S, Prunier-Leterme N, Le Trionnaire G (2022) Masculinization of the X-chromosome in aphid soma and gonads. bioRxiv, 2021.08.13.453080, ver. 4 peer-reviewed and recommended by Peer Community in Evolutionary Biology. https://doi.org/10.1101/2021.08.13.453080.

## Abstract

Males and females share essentially the same genome but differ in their optimal values for many phenotypic traits, which can result in intra-locus conflict between the sexes. Aphids display XX/X0 sex chromosomes and combine unusual X chromosome inheritance with cyclical parthenogenesis. Theoretical and empirical works support the hypothesis that the large excess of male-biased genes observed on the aphid X chromosome compared to autosomes evolved in response to sexual conflicts, by restricting the products of sexually antagonistic alleles to the sex they benefits. However, whether such masculinization of the X affects all tissues (as expected if it evolved in response to sexual conflicts) or is limited to specific tissues remains an open question. Here, we measured gene expression in three different somatic and gonadic tissues of males, sexual females and parthenogenetic females of the pea aphid. We observed a masculinization of the X in each of the studied tissues, with male-biased genes being 2.5 to 3.5 more frequent on the X than expected. We also tested the hypothesis that gene duplication can facilitate the attenuation of conflicts by allowing gene copies to neo- or sub-functionalize and reach sex-specific optima. As predicted, X-linked copies of duplicated genes having their other copies on autosomes were more frequently male-biased (40.5% of the genes) than duplicated autosomal genes (6.6%) or X-linked single-copy genes (32.5%). These results highlight a peculiar pattern of expression of X-linked genes in aphids at the tissue level and provide further support for sex-biased expression as a mechanism to attenuate intra-locus sexual conflicts.

## Introduction

Sexual dimorphism, the difference between males and females at any phenotypic trait such as behavior, morphology, physiology or life history, is widespread. These differences are pervasive among eukaryotes, from plants to nematodes, insects, birds and mammals, to name a few (Cox and Calsbeek 2009; Williams and Carroll 2009). Regardless of its extent, sexual dimorphism engages males and females in a constant tug-of-war because their reproductive interests (such as optimal mating rate, number of partners, parental investment…) never align, owing to constitutive investment differences in gametes and/or progeny (Bonduriansky and Chenoweth 2009).

Differences in optimal trait values between sexes may generate intra-locus sexual conflicts. Typically, a new allelic variant could be beneficial to a female but deleterious to a male or *vice versa*. Such a sexually antagonistic (SA) allele is predicted to increase in frequency as long as the cost/benefit balance is positive. This increase leads to a so-called gender load in the population, due to the transmission of SA alleles to both sons and daughters (Chippindale et al. 2001; Rice and Chippindale 2002; Bonduriansky and Chenoweth 2009).

Several mechanisms may alleviate gender load (Bonduriansky and Chenoweth 2009). One is the evolution of sex-biased or sex-specific gene expression through a modifier of expression (Rice 1984). Once a SA allele is frequent enough, the reduction of its expression in the sex where it is deleterious may allow this variant to further increase in frequency and to possibly reach fixation (Rice 1984; Ellegren and Parsch 2007; Bonduriansky and Chenoweth 2009). This implies that the reduction of expression of the SA allele is beneficial to individuals of this sex. For genes that must be expressed at a certain level, a gene duplication event could allow bringing a new gene copy to sub- or neo-functionalize toward the sex-specific optimum (Bonduriansky and Chenoweth 2009; Connallon and Clark 2011; Gallach and Betrán 2011). Interestingly, these two processes (the duplication and the change in expression) could occur simultaneously, when the duplicated copy inserts in a region of the genome that already shows specific expression pattern (e.g., Arthur et al. 2014).

The invasion of the population by a SA allele and the attenuation of gender load through duplication and/or evolution of sex-biased gene expression may take place at different timescales. Indeed, the increase in frequency of a SA allele can be as rapid as a few generations, depending on its effect on fitness (e.g., Dean et al. 2012 for an experimental demonstration). The attenuation of the conflict by expression change or gene duplication may take much longer as it relies on rare random events, themselves depending on effective population size and mutation rate (Rice 1984; Stewart et al. 2010; Connallon and Clark 2011; Collet et al. 2016).

Importantly, the conditions for invasion by a SA allele differ between autosomes and sex chromosomes (Rice 1984; Fry 2010). In XX/XY systems, any SA allele that benefits males can invade the Y without conflict assuming complete linkage between the SDR (sex-determining region) and the SA locus. The picture for the X is more complex (Vicoso and Charlesworth 2006). X-linked recessive alleles are exposed to selection in males, while the female-biased transmission of the X (X chromosomes are transmitted twice more often by females than by males) gives more importance to selection episodes occurring in females. As a result, the X should accumulate recessive male-beneficial alleles and dominant female-beneficial ones. Similar processes are expected to occur in ZZ/ZW systems (e.g., birds, lepidopterans…).

Aphids constitute an interesting model to study the evolution of SA alleles as they show an XX/X0 sex-determining system combined with cyclical parthenogenesis: the alternation between several parthenogenetic generations in spring and summer and a single sexual generation in autumn. As a result, three distinct reproductive morphs occur in aphids: males, sexual females and parthenogenetic (asexual) females. Sexual females are genetically identical to their parthenogenetic mother, while male production involves the random elimination of one of the X (Wilson et al. 1997). Furthermore, during spermatogenesis only sperm cells carrying an X chromosome develop (Blackman 1987), so that the fusion of a sperm cell (AX) and an ovum (AX) always produces a diploid individual at the X and autosomes, which develops into a parthenogenetic female.

Theoretical models (Jaquiéry et al. 2013) predict that the peculiar inheritance of the X in aphids, the alternation between sexual and asexual reproduction, and the presence of three different morphs (sexual females, parthenogenetic females and males) have a major influence on the genomic location of SA allelic variants. In particular, conditions for the invasion of variants that are beneficial to males and deleterious to parthenogenetic females are predicted to be less restrictive for the X than for autosomes. By contrast, the conditions for the invasion of variants that are detrimental to males and beneficial to parthenogenetic females are more restrictive for the X. These models thus predict the X to be optimized for male functions. Genomic analyses on the pea aphid *Acyrthosiphon pisum* showed that the X chromosome had a large excess of genes preferentially expressed in males (i.e., male-biased genes) compared to autosomes, and a deficit of parthenogenetic female-biased genes, resulting in a “masculinization” of this chromosome (Jaquiéry et al. 2013). This pattern matched predictions made under the hypothesis that evolution of sex-biased gene expression reduces sexual conflicts by decreasing the expression of a sexually antagonistic allele to the sex it benefits (Rice 1984). Interestingly, masculinization of the X has also been observed in another aphid species (*Myzus persicae*) that diverged from the pea aphid lineage 40 MYA (million years ago) (Mathers et al. 2019), but not in psyllids (Li et al. 2020) – a closely related group to aphids but which undergoes obligate sexual reproduction. These studies provide further support that the masculinization of the X evolved in response to intra-locus sexual conflicts resulting from the peculiar life cycle (cyclical parthenogenesis) and X inheritance in aphids. However, as previous studies on aphids analyzed whole-body transcriptomes (Jaquiéry et al. 2013; Jaquiéry et al. 2018; Mathers et al. 2019; Li et al. 2020), it is unknown whether the observed masculinization of the X systematically occurs within each type of tissue or is driven by some specific tissue with unusual expression patterns. Indeed, empirical studies in model species revealed considerable variation between tissues. For example, other factors such as meiotic-sex chromosome inactivation (MSCI) makes sex-chromosomes an inappropriate location for spermatogenesis genes in *Drosophila* and mammals (e.g. Khil et al. 2004, Vibranovski et al. 2009). Dosage compensation may also differ between tissues (Nozawa et al. 2013, Vensko and Stone 2015), which would explain why the *Drosophila* X is enriched in male-biased genes expressed in the brain, but shows no excess or even a deficit of male-biased genes for all other tissues (Huylmans et al. 2015). Given the scarce knowledge of dosage compensation and MSCI in aphids (the only four studies that analyzed gene expression in aphid males were performed on whole individuals, Jaquiéry et al. 2013, Mathers et al. 2019, Liu et al. 2021, Ziabari et al. 2022), we were not able to account for these factors in our theoretical predictions. Thus, we focused primarily on testing predictions assuming sexual antagonism, but explored the presence of dosage compensation (and MSCI to a lesser extent) on the tissue-specific transcriptomes collected for this study.

Here, we predicted that if intra-locus sexual conflict is a strong driver of the masculinization of the aphid X chromosome, masculinization would occur in all tissues. To verify this prediction, we measured gene expression in different tissues from males, sexual females and parthenogenetic females, including gonadic and somatic tissues. Sex-biased genes were more frequent in gonads than in less sexually dimorphic tissues, nevertheless we observed a masculinization of the X in each type of tissue, suggesting that - in each tissue - male-beneficial alleles were favored on the X and that intra-locus conflict may be resolved through the evolution of sex-biased gene expression. Moreover, we confirmed that the X-linked copy of a duplicated gene having another copy on autosomes is more likely to show a male-biased expression than its autosomal copy or an X-linked single-copy gene. This result suggests that duplications facilitate sub- or neo-functionalization toward the sex-specific optimum.

## Results

### Gene expression levels

Gene expression levels in three different tissues (heads, legs and gonads) of the three morphs (males, sexual females and parthenogenetic females) were measured from RNA-seq counts on individuals produced by the same pea aphid clonal lineage (supplementary table S1). Overall, 14,605 genes out of the 20,639 predicted genes were expressed (> 1 count per million reads [CPM] in at least two samples) in the 18 samples (3 morphs × 3 tissues × 2 replicates). We assigned 18,719 (90.7%) of the 20,639 predicted genes as autosomal or X-linked, based on scaffold assignments from Jaquiéry et al. (2018). The genes that were not assigned (9.3%) were located on scaffolds or part of scaffolds that were not clearly assigned to X or autosomes in Jaquiéry et al. (2018) and they were thus not considered in the subsequent analyses. Only 51% (3044/5961) of the X-linked genes were found to be expressed, against 85% (10,890/12,758) of the autosomal genes (figure 1A). The genes identified as not expressed in the 18 samples also generally showed no or low expression in whole-body RNAseq of males and females (Jaquiéry et al. 2013), especially for X-linked genes (see supplementary text S1 for details). On average, more genes were expressed in the samples from the different male tissues than in female samples, especially for X-linked genes (X-chromosome: median number of expressed genes in males = 2020, median in females = 1425, two-sided Mann-Whitney test, *p* = 0.0009; autosomes: median in males = 9061, median in females = 8560, two-sided Mann-Whitney test, *p* = 0.067, figure 1B).

**Figure 1:**
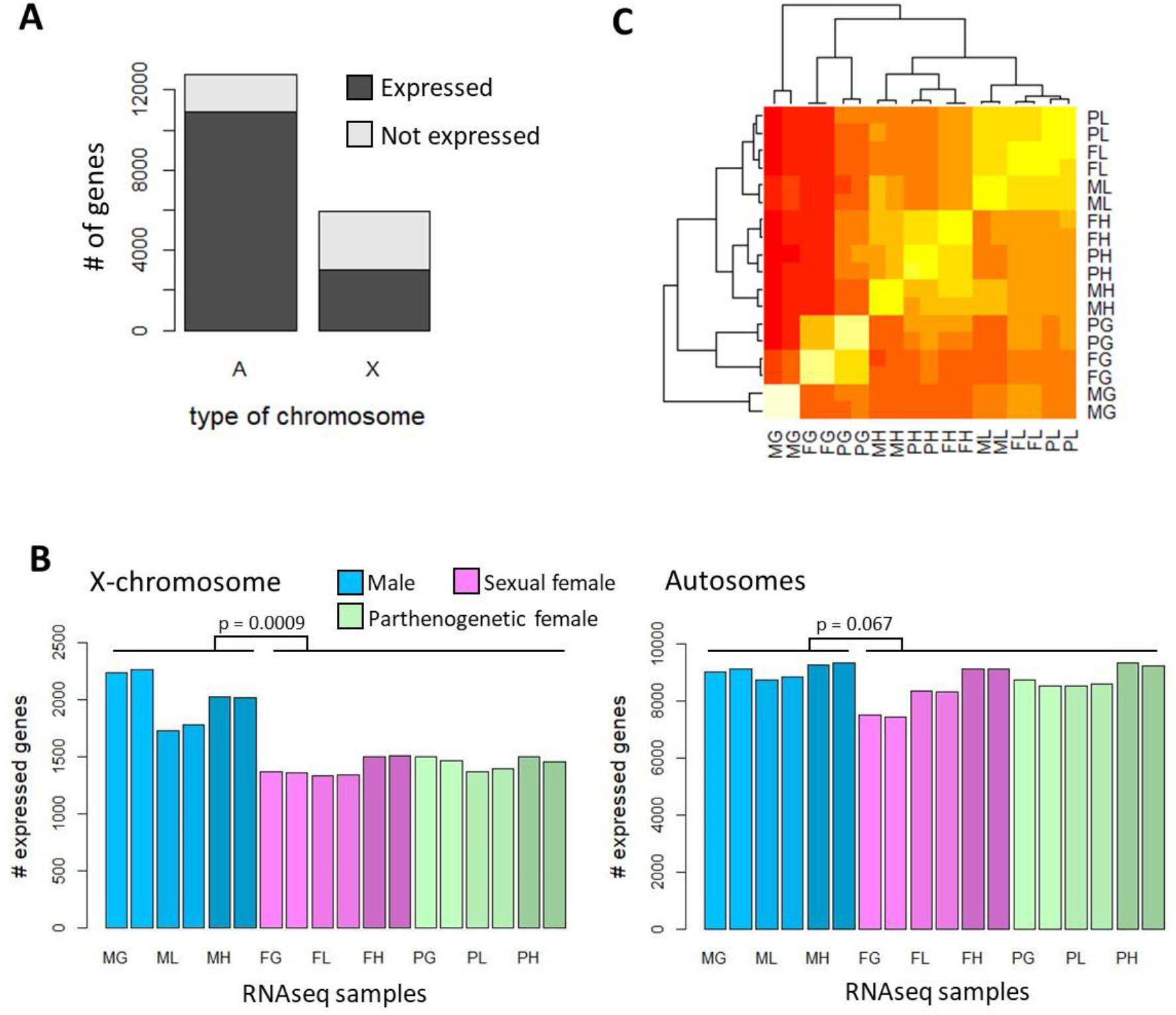
Gene expression in the 18 different RNA-seq samples. A) Number of genes considered as expressed (CPM>1 in at least two samples) and not expressed on the X and on the autosomes. B) Number of expressed genes per sample (expressed at more than 1 CPM) for X-linked and autosomal genes (significance of differences between the X and autosomes was estimated with two-sided Mann-Whitney tests). C) Heatmap of log(CPM+1). Samples group by tissue for head (H) and leg (L) samples, with the parthenogenetic (P) and sexual female (F) samples being always more similar compared to the male samples (M). Expression patterns of male gonad samples (MG) are the most divergent. MG: male gonad, MH: male head, ML: male leg, FG: sexual female gonad, FH: sexual female head, FL: sexual female leg, PG: parthenogenetic female gonad, PH: parthenogenetic female head, PL: parthenogenetic female leg.

In the heatmap based on gene expression levels (figure 1C, supplementary figure S1), samples grouped systematically by replicate of the same condition, and then by tissue for leg and head samples. Within each of these tissues, the four female samples were always more similar to each other than to the male samples. Gonad samples were the most heterogeneous ones, samples from testes (MG) being highly different from all other samples, and samples from parthenogenetic and sexual female gonads grouping together.

### The X chromosome is enriched in male-biased genes at the tissue-level

To test whether some masculinization of the X was observed at the tissue-level, we first categorized genes according to their relative expression patterns in the different conditions. We defined a gene as “biased” toward, or preferentially expressed in, a set of samples (which can be a particular tissue from a particular morph, all the tissues from a particular morph or a particular tissue in all morphs) when at least 70% of all reads mapping to this gene were observed in this set of samples (see methods). Note that increasing this threshold to 80% or 90% or decreasing it to 60% or 50% did not qualitatively change the results (supplementary figure S2). Testes showed the highest number of biased genes, with 1170 genes (referred to as MG+ genes) being preferentially expressed in this tissue. Then came sexual female ovaries, with 375 FG+ genes, and male heads, with 203 MH+ genes (figure 2). Overall, the number of sex-biased genes was higher in highly sexually dimorphic tissues (gonads) than in less sexually dimorphic tissues (heads and legs). Tissue-biased genes (i.e., genes expressed mainly in a tissue of all morphs) were common. Heads showed the highest number of tissue-biased genes (1169 H+ genes), followed by legs (607 L+ genes) and gonads (511 G+ genes). Contrastingly, morph-biased genes (i.e., genes expressed mainly in a morph in all tissues) were much less frequent for females (only 58 F+ and 130 P+ genes) than for males (596 M+ genes) (figure 2). When considering all genes preferentially expressed in a given morph, without considering tissues (e.g., by summing MG+, MH+, ML+ and M+ genes for males), a total of 2001, 501 and 299 genes were biased toward males, sexual females and parthenogenetic females, respectively.

**Figure 2:**
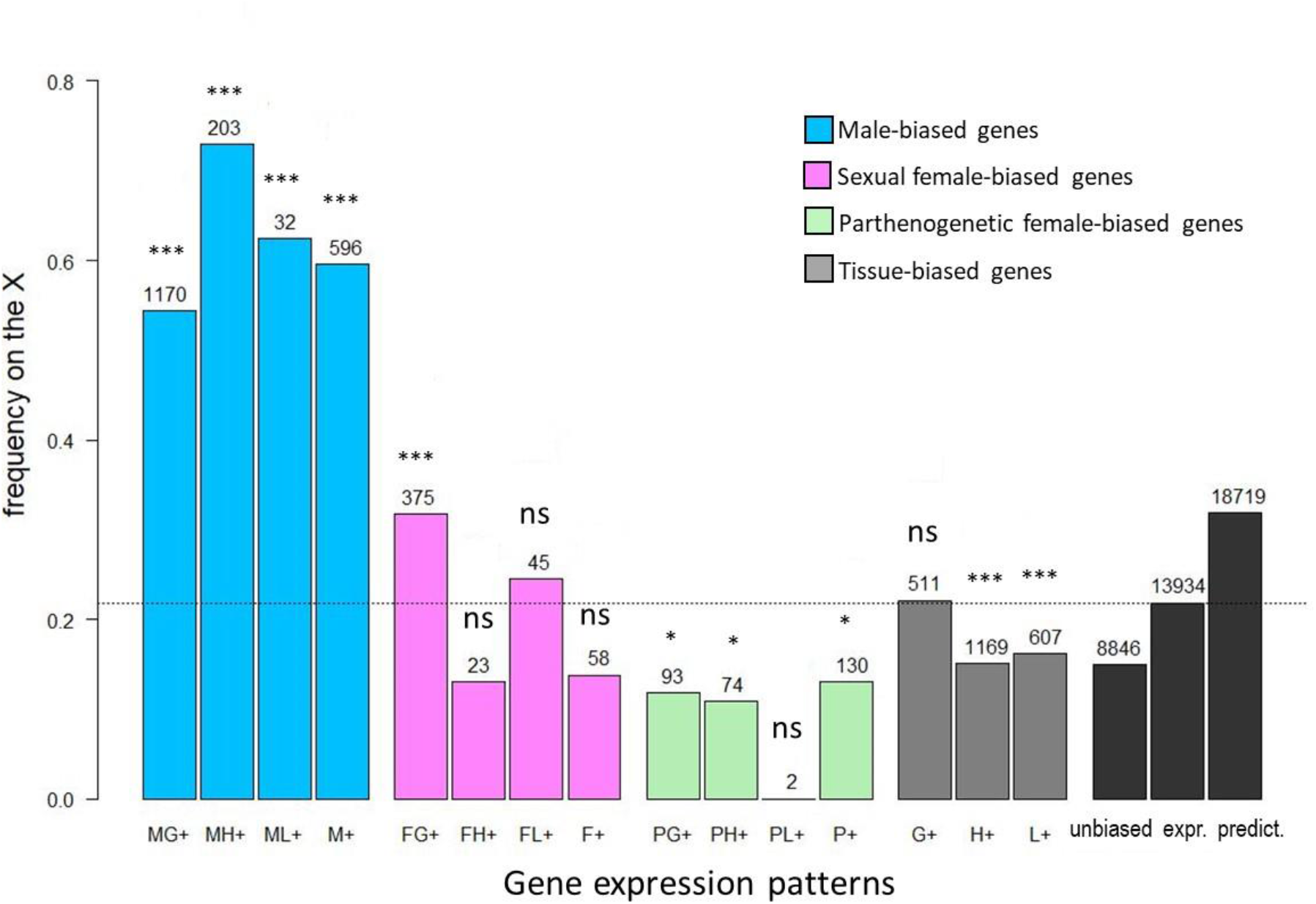
Proportions of X-linked genes among genes preferentially expressed in various morphs and/or tissues. Blue bars represent male-biased genes; MG+, MH+ and ML+: genes expressed preferentially in male gonads, heads and legs, respectively. M+: genes preferentially expressed in males when pooling all tissues,excluding the genes assigned to the previous categories. Pink bars represent sexual female-biased genes, with F standing for females and letters G, H and L having the same meaning as in males. Green bars represent parthenogenetic female-biased genes (P). Grey bars represent genes expressed preferentially in one of the tissues (gonads, heads or legs) and not limited to a particular morph. Black bars represent the frequency of X-linked genes among genes with unbiased expression (“unbiased”), genes expressed with CPM > 1 in at least two libraries (“expr.”) or all predicted genes (“predict.”). The horizontal dotted line represents the proportion of X-linked gene among expressed genes. The number of genes from each category is shown above bars, as well as the p-value (two-sided binomial tests against the expected frequency on the X chromosome estimated from expressed genes, which corresponds to the dotted horizontal black line). ***: *p* < 0.001; **: *p* < 0.01; *: *p* < 0.05; ns: *p* ≥ 0.05.

Interestingly, these different categories of genes differed in their chromosomal locations (figure 2, supplementary table S2). The proportions of X-linked genes among genes expressed preferentially in testes (MG+), male heads (MH+), male legs (ML+) or simply in males regardless of tissue (M+) varied from 54% to 73%, and significantly exceeded (two-sided binomial tests, *p* < 10^−7^ in all cases) the null expectation, which we took as the proportion of X-linked genes among all expressed genes (22%) (if we consider all predicted genes – supported by expression data or not – 31.8% locate on the X). Contrastingly, genes that were preferentially expressed in parthenogenetic females were less likely to locate on the X than expected (two-sided binomial tests, *p* ranging from 0.014 to 0.023 for PG+, PH+ and P+, not significant for PL+), with proportions of X-linked genes ranging from 0% to 13% depending on tissues. The proportion of X-linked genes among sexual female-biased genes were intermediate (13% to 32%), with only those preferentially expressed in sexual female gonads being more frequent on the X (two-sided binomial test, *p* = 10^−5^). Genes that were preferentially expressed in gonads (G+) showed no deviation from the null expectation, as 22% of them located on the X, while genes preferentially expressed in heads (H+) and in legs (L+) were significantly less frequent on the X (15% to 16%) than expected (two-sided binomial tests, *p* = 10^−8^ and *p* = 0.0006, respectively). Baring the strong difference between the X and autosomes, the distribution of biased genes within chromosomes was rather homogeneous (supplementary figure S3).

We found that the breadth of gene expression, measured with τ (an index that ranges from of 0 – indicating similar expression in all conditions – to 1 – indicating expression in one condition only) was significantly narrower for X-linked genes than for those on autosomes (median *τ_x_*= 0.71, *τ_A_*= 0.33, Mann-Whitney U-test, U = 23,201,000, *p* < 10^−15^).

### Expression patterns of single- and two-copy genes

To investigate the extent to which gene duplication facilitates the evolution of gene expression toward the sex-specific optimum, we compared the expression of autosomal and X-linked genes that belong to single-copy and multicopy gene families. Multigenic gene families were identified by Boulain et al. (2018) from orthoDB on 17 arthropod genomes.

We found that the X chromosome contained more genes that belong to multicopy families than autosomes (38% of the genes on the X belong to multicopy families, against 28% for autosomal genes). When restricting our analyses to genes supported by expression data, 1633 and 7428 single-copy genes locate on the X and on autosomes, respectively. We also found 210 gene families composed of two expressed genes with one being on the X and the other on autosomes. On autosomes, the percentages of genes with male-biased expression (combining M+, MG+, ML+ and MH+ genes) were very similar between single- and two-copy genes, at 6.5% and 6.6% respectively (figure 3, Chi-squared test, χ^2^ ≈ 0, df = 1, *p* = 1). On the X chromosome however, the proportion of male-biased genes was significantly higher for two-copy genes (40.5%) than for single-copy genes (32.5%) (Chi-squared test, χ^2^ = 5, df = 1, *p* = 0.025), these two proportions being much higher than their equivalents on autosomes (Chi-squared tests, single-copy genes: χ^2^ = 939.5, df = 1, *p* < 10^−15^, two-copy genes: χ^2^ = 64.8, df = 1, *p* < 10^−15^). Sexual female- and parthenogenetic female-biased genes accounted only for a few percent of single- and two-copy genes. These female-biased genes showed minor differences in proportion between chromosomes (significant for single-copy genes only for parthenogenetic female [χ^2^ = 4.9, df = 1, *p* = 0.027] and for sexual female [χ^2^ = 8.9, df = 1, *p* = 0.003], figure 3). They constituted similar proportions of the single- and two-copy genes on a given chromosome type (supplementary table S3).

**Figure 3:**
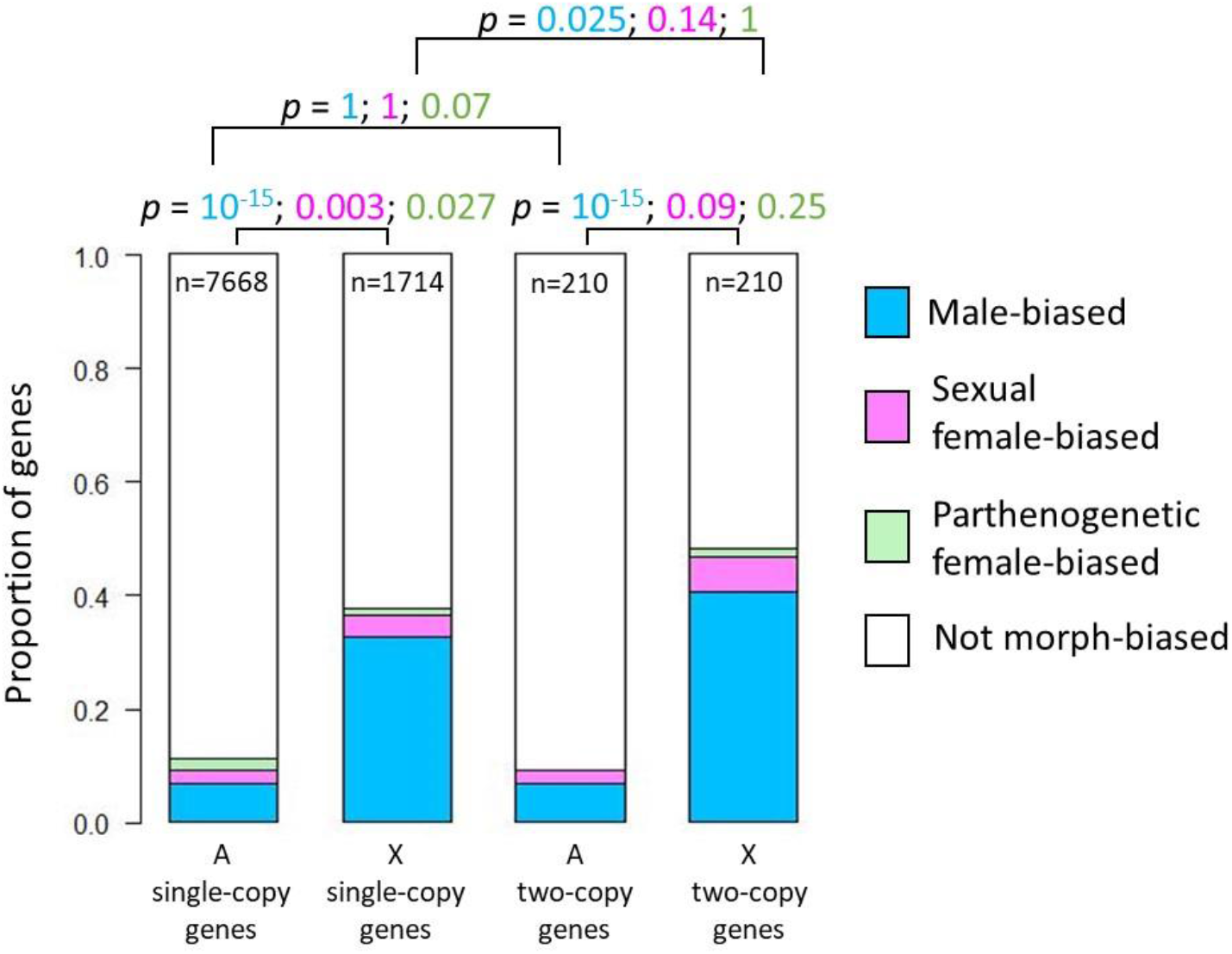
Proportion of genes showing preferential expression in different morphs, according to their number of copies (single or two copies) and chromosomal location (A: autosomes, X: X chromosome). Each two-copy gene has one copy on an autosome and the other on the X. P-values (Chi-squared tests) are shown, with font colors corresponding to the tested morph, according to the color of sectors. The number of genes composing each distribution is indicated on the plots.

### Expression levels in morphs and tissues

The median expression levels of X-linked genes in somatic tissues and gonads from male and sexual female morphs were systematically lower than those of autosomal genes (figure 4, *p* < 10^−15^ in all comparisons, two-sided Mann-Whitney tests), irrespective of the dose of X chromosomes per cell (two for sexual females and one for males). The same patterns were observed in parthenogenetic females (*p* < 10^−15^ in all comparisons, supplementary figure S4). The mode of Log2 ratio of male-to-female RPKM (using sexual females in figure 4CFI and parthenogenetic females in supplementary figure S4) lies close to 0 for both autosomal and X-linked genes, indicating dosage compensation for non-sex-biased genes in gonads and somatic tissues. Yet, we observed an excess of genes with high Log2 ratio of male to female expression, especially for the X chromosome in gonads and heads (figure 4CF). This indicates an overexpression of some of the genes located on the single X chromosome of male cells, which exceeds dosage compensation. This pattern was expected, given that male-biased genes are significantly more frequent on the X than on autosomes (figure 2).

**Figure 4:**
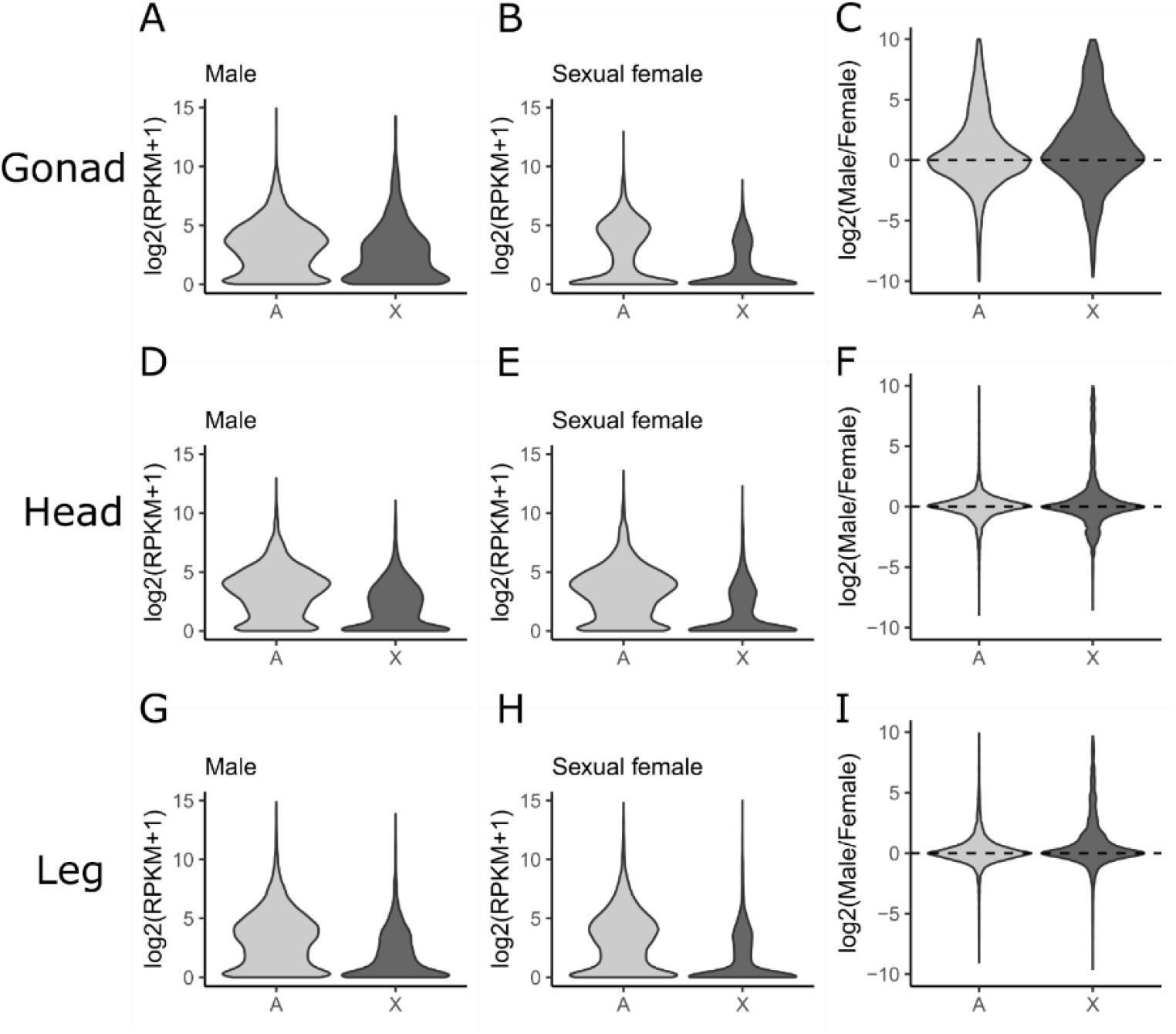
Logarithm of gene expression (RPKM, reads per kilobase per million mapped reads) for X-linked and autosomal genes for the different types of libraries (gonads, heads or legs of males and sexual females). X-linked genes are significantly less expressed than autosomal genes in all cases (two-sided Mann-Whitney tests, *p* < 10^−15^). Logarithm of male-to-sexual female ratio of RPKM is also shown for the three different tissues.

When we removed the genes characterized by a fold-change larger than 2 between males and sexual females in at least one of the tissues (i.e. sex-biased genes), we still found strong evidence for dosage compensation in the three tissues (supplementary figure S5). We also observed that dosage compensation occurs throughout the X chromosome for all 3 tissues (supplementary figure S6BDF), although the terminal portion of this chromosome appeared to be particularly rich in sex-biased genes (supplementary figure S6ACE and S3).

## Discussion

Because of the peculiar inheritance of the X chromosome in aphids – males transmit systematically their unique X to all sperm cells leading to the production of female-only progeny –, the presence of three distinct morphs (sexual females, parthenogenetic females and males) and the alternation between sexual and asexual reproduction, a specialization of the X into male functions is expected (Jaquiéry et al. 2013). Indeed, models have shown that the conditions for invasion by male-beneficial/parthenogenetic female-detrimental SA alleles are less restrictive for the X than for autosomes, while the opposite is true for male-detrimental/parthenogenetic female-beneficial alleles. SA alleles that are favorable to sexual females should show little bias, the direction of which (i.e., the depletion or enrichment of the X with genes carrying such variants) depending on the selective effect of the allele on males and parthenogenetic females (table 1). A key finding of these predictions is that they are not qualitatively affected by allele dominance level *h* (the aphid X chromosome is a preferred location for male-beneficial alleles for all values of *h*≠1, see Jaquiéry et al. 2013). This contrasts with other X0 and XY species, where the X accumulates both recessive (*h*<0.5) male-beneficial alleles and dominant (*h*>0.5) female-beneficial alleles (Vicoso and Charlesworth 2006; Ellegren and Parsch 2007). Consequently, simpler predictions can be made on aphids: the X should be enriched with male-beneficial alleles, parthenogenetic female-beneficial alleles should be more common on autosomes (being counter selected on the X), and sexual-female beneficial alleles should show no consistent bias (table 1).

**Table 1:** Preferred chromosomal location (X chromosome *versus* autosomes) of different types of alleles with morph-antagonistic effects (*s_m_*, *sf* and *s_p_* stand for the selective coefficient of the new variant on males, sexual females and parthenogenetic females, respectively). Predictions originate from analytical and simulation models developed in Jaquiéry et al. (2013).

| Fitness effect of a SA mutation | Preferred chromosomal location in aphids |
| --- | --- |
| $s_m > 0, s_f < 0, s_p < 0$ | Favored if on the X |
| $s_m > 0, s_f > 0, s_p < 0$ | Favored if on the X |
| $s_m < 0, s_f < 0, s_p > 0$ | Disfavored if on the X |
| $s_m < 0, s_f > 0, s_p > 0$ | Disfavored if on the X |
| $s_m < 0, s_f > 0, s_p < 0$ | Slightly disfavored if on the X |
| $s_m > 0, s_f < 0, s_p > 0$ | Slightly favored if on the X |

Although the development of high-throughput approaches have allowed to pinpoint putative SA genes (Innocenti and Morrow 2010; Lucotte et al. 2016; Ruzicka et al. 2019), identifying these genes and estimating sex-specific selection and dominance coefficients are challenging tasks that have been achieved in only a handful of studies (Barson et al. 2015; Husby et al. 2015). However, since the evolution of a lower expression level in the sex that suffers from an antagonistic allele could help resolving intra-locus sexual conflicts and allow the SA allele to reach fixation (Rice 1984; Vicoso and Charlesworth 2006; Ellegren and Parsch 2007), the chromosomal location of sex-biased genes can be used to indirectly test predictions.

Here, we analyzed gene expression in different tissues of distinct morphs in the pea aphid and found that the chromosomal location of morph-biased genes followed the predictions made under SA models from Jaquiéry et al. (2013). In the three tissues considered, male-biased genes were largely overrepresented on the X, parthenogenetic female-biased genes were underrepresented, and sexual female-biased genes showed no consistent bias. These empirical data thus support the hypothesis that, in aphids, sexual conflicts would be the key driver of the masculinization of the X and of the specialization of autosomes for the parthenogenetic phase of the life cycle.

Genes showing a male-biased expression were much more frequent on the X chromosome than expected under a random distribution, irrespective of the tissue considered. Genes expressed mainly in testes, male heads and male legs were 2.5 to 3.5 times more frequent than expected on the X, so were genes that were male-biased in all the tissues considered. As all the investigated tissues contribute to this pattern, these results extend previous studies reporting a general enrichment in male-biased genes (measured from whole-body samples) on the aphid X chromosome (Jaquiéry et al. 2013; Jaquiéry et al. 2018; Mathers et al. 2019). Overall, this and previous studies demonstrate strong and consistent bias toward the X in aphids for genes expressed predominantly in males. Interestingly, on the X chromosome, the percentage of male-biased genes was significantly higher for two-copy genes (40.5%) than for single-copy genes (32.5%), while these percentages were similar for autosomal genes (6.5% and 6.6%). These results match the predictions that duplications may facilitate sub- or neo-functionalization toward sex-specific optima, and that, for genes located on the X chromosome, the male optimal expression levels are favored rather than the female optima.

Conversely, parthenogenetic female-biased genes were significantly less frequent on the X, except for parthenogenetic female leg-biased (PL+) genes due to lack of power, as only two such genes were found, both located on autosomes. This matches previous observations on whole body transcriptomes (Jaquiéry et al. 2013; Mathers et al. 2019). This consistent pattern therefore reveals a reduction of the contribution of the X chromosome to biological functions and processes occurring in parthenogenetic females, and corroborates the observation that the X is depleted from functionally important genes for the parthenogenetic phase (Li et al. 2020). Interestingly, the chromosomal location of sexual female-biased genes did not significantly depart from random expectations, except for genes specifically expressed in ovaries. These were significantly more frequent than expected on the X, although their proportion (32%, figure 2) was much lower than for male-biased genes (54%−73%). According to the models (table 1), variants of X-linked genes that are beneficial to sexual females should increase in frequency only if they are also beneficial to males. This could be the case for genes controlling sexual reproduction-related functions that were not sex-specific (e.g., meiosis). In this case however, we do not expect such genes to be preferentially expressed in sexual females. These genes may be less/not expressed in adult males because spermatogenesis is often already completed at this stage (Wieczorek et al. 2019). Alternatively, it is possible that these genes had variants that were beneficial to both sexes, which have since evolved functions that are more specific to sexual females.

Among hemipterans, the accumulation of male-biased genes on the X appears to be specific of aphids. A deficit of male-biased genes on the X was observed in three hemipteran species (two heteropteran bugs and a leafhopper) that only reproduce sexually and are very distantly related to aphids, despite an apparent homologous origin of the X chromosome (Pal and Vicoso 2015; Mathers et al. 2021). In psyllids, which diverged more recently (~200–250 MYA) from aphids and are characterized by obligate sexual reproduction and X0 sex determination, no enrichment of male-biased genes on the X chromosome was found (Li et al. 2020). Hence, the accumulation of male-biased genes on the X chromosome in aphids would have started after the divergence between aphids and psyllids. This scenario supports a role for cyclical parthenogenesis (which appeared ~200 MYA in aphid ancestors, Davis 2012) in the masculinization of the X in aphids.

While the chromosomal location of the different types of sex-biased genes matches the predicted evolution of gene expression as a mitigation mechanism of sexual conflicts, other processes could contribute to the observed patterns. Genes located on different chromosomes types could differ in other characteristics that could result in a non-random genomic distribution of male-biased genes. For example, the X chromosome of *Drosophila* is enriched in young genes (those that are present in only a restricted taxonomic group), which are more likely to show sex-biased expression (Palmieri et al. 2014). In our analysis however, the X-linked copy of a two-copy gene had a much greater probability of being male-biased (0.405) than its autosomal counterpart (0.066, figure 3), demonstrating that the masculinization of the X does not solely reflect differences in gene characteristics. Moreover, the three *A. pisum* autosomes are systematically depleted in male-biased genes and enriched in parthenogenetic female-biased genes (supplementary table S4, supplementary figure S3). This reinforces the specific hemizygosity of the X in males as a determinant factor for the observed patterns. Consequently, we do not see an alternative hypothesis to explain the accumulation of male-biased genes on the X to the one based on the resolution of sexual conflicts by an evolution of sex-specific or sex-biased gene expression.

The number of morph-biased genes was highest for gonadic tissues, with 1170, 375 and 130 genes being specific of gonads of males, sexual females or parthenogenetic females, respectively. This observation likely reflects the highly morph-specific functions of gonads: the production of sperm in males, of yolky eggs in sexual females and of embryos in viviparous parthenogenetic females (Michalik et al. 2013). In other species, testis also stands out as the tissue showing the most specific gene expression patterns (Meiklejohn and Presgraves 2012; Uhlén et al. 2015). Many genes also showed preferential expression in male heads (202), against 23 and 93 for sexual and parthenogenetic female heads, respectively. This could reflect sensorial and/or behavioral differences between males and females: males have to actively search for females and initiate mating, while females spend most of their time in feeding. Legs showed a low number of morph-biased genes, most of which were overexpressed in sexual females (45 genes, against 32 for males and 2 for parthenogenetic females). This result may reflect the existence of specific organs on sexual female tibias (scent plaques), which are responsible for the secretion and release of sex pheromones (Murano et al. 2018). Overall, the degree of sex-biased expression varied with the degree of specialization of each tissue in sexual functions. Whether this suggests that sexual antagonism is stronger in such highly sexually dimorphic tissue than in tissues with more similar function in both sexes remains an open question, as causal demonstration of the link between sex-biased expression and sex-specific fitness is challenging (Mank 2017).

Interestingly, the strong enrichment of the X with male-biased genes is similar across tissues (ranging from 54% to 73%). This result strikingly contrasts with *Drosophila*, where male-biased genes from different tissues show opposing patterns: male-biased genes in brains are strongly enriched on the X, while there is either no departure from random expectation or a paucity of male-biased genes on the X for the other tissues (Huylmans and Parsch 2015). These differences were interpreted as resulting from the interplay between dosage compensation (the brain could be more sensible to gene dose, and thus would require a tighter dosage control) and sex-specific regulation of gene expression (Huylmans and Parsch 2015). In aphids, the consistent enrichment of the X with male-biased genes in different tissues suggests that similar evolutionary forces apply to solve SA conflicts in reproductive and somatic tissues. Under the hypothesis that sex-biased expression evolved in response to sexual conflicts, our results suggest that their attenuation occurs in all tissues. Our data also indicate that - even if sexual conflicts could be more frequent in gonads (assuming that sex-biased genes reveal past or ongoing conflicts) - conflicts also occur in a wide range of somatic tissues. This pattern was also observed by Innocenti and Morrow (2010), who found that transcripts showing signature of sexual antagonism in *Drosophila* were frequent in soma.

The high expression of a substantial number of X-linked genes in males despite the haploid state of this chromosome in this sex is intriguing. The X chromosome shows several specific characteristics, among which a slightly larger amount of intra-chromosome duplicates (449 for 132 Mb) compared to the largest autosomes (413 for 170 Mb, Li et al. 2019), and an enrichment with multi-copy orthologs (Li et al. 2020). Our analyses confirmed this trend, as 38% of the genes on the X belongs to multicopy families, against 28% for autosomal genes. It is yet to be determined whether recent (undetected) duplications are more common on X and could contribute to high expression of some of the X-linked genes in males. However, epigenetic mechanisms probably play a more important role. Indeed, the X-chromosome is hypermethylated in male aphids compared to autosomes, and differential gene methylation between males and females positively correlates with differential expression, especially for the X chromosome (Mathers et al. 2019). An increased accessibility of the chromatin of the X in males was also documented (Richard et al. 2017).

Another interesting feature of the aphid X chromosome is its larger fraction of unexpressed genes compared to autosomes, amounting to half of the X-linked genes based on our tissue samples data (against 15% of the autosomal genes). This characteristic was also underlined from whole body transcriptomes (Jaquiéry et al. 2013; 2018; Richard et al. 2017; Li et al. 2020; Mathers et al. 2019). Two factors may contribute to the large number of X-linked genes classified as unexpressed. X-linked genes might show a narrower expression breadth, being restricted to some (unsampled) morphs, tissues or stages. The exceptionally high τ value for expressed X-linked genes indeed provides some support to this hypothesis. Nevertheless, a large fraction of the X-linked genes classified as “unexpressed” here also shows low expression support in whole body libraries (see supplementary text S1): 40% are supported by 0 reads and 34% by 1-5 reads. It is thus probable that both effects (absence of expression or narrower expression breadth) explain the large fraction of unexpressed X-linked genes. RNA sequencing on a larger diversity of tissues (especially in males) and stages may be required to resolve this point. Interestingly, Li et al. (2020) found that genes considered as “functionally important” were less likely to locate on the X chromosome. The evolutionary forces that drive these patterns remain to be identified, but they could be linked to sexual antagonism or to the particular epigenetic state of the X.

Previous studies suggested dosage compensation in the pea aphid and the green peach aphid from whole-body transcriptomes (Jaquiéry et al. 2013; Pal and Vicoso 2015; Richard et al. 2017; Li et al. 2020; Mathers et al. 2019). Here, we show dosage compensation in all investigated tissues, including testis. While single-cell transcriptomics will be essential to demonstrate or refute that X-linked genes are dosage compensated in various cell types during spermatogenesis, dosage compensation in testis would be another peculiarity of aphids (but see Witt et al. 2021; Mahadevaraju et al. 2021). Indeed, sex chromosomes of other dosage-compensated species seem generally not compensated in the gonads of the heterogametic sex in diptera and lepidoptera, at the scale of entire organs (Vicoso and Bachtrog 2015; Gu et al. 2017; Gu and Walters 2017). However, single-cell RNAseq analysis of *Drosophila* testis has evidenced dosage compensation in pre-meiotic and somatic testis cells (Witt et al. 2021, Mahadevaraju et al. 2021). In other groups (mammals, birds, nematodes, fungi), sex-linked genes are silenced by meiotic sex chromosome inactivation or MSCI (Shiu et al. 2001; Bean et al. 2004; Turner 2007; Schoenmakers et al. 2009; but see Guioli et al. 2012; Daish et al. 2015), and recent studies suggest that MSCI could also occur in *Drosophila* (Mahadevaraju et al. 2021; Witt et al. 2021). Different hypotheses have been proposed to explain the silencing of sex-linked genes during gametogenesis. MSCI could be a consequence of the mechanisms that protect against unwanted recombination between X and Y and allow DNA repair in its absence (Lu et al. 2015). MSCI may also has evolved as a mean to protect unsynapsed chromosomes from the invasion of transposons (Huynh et al. 2005) or segregation distorters that would bias sex-ratio (Meiklejohn and Tao 2010). If protection from segregation distorters is an important driver of sex-linked gene silencing, this could explain why the expression of X-linked genes in aphid testis has not been repressed during evolution and thus can be subject to dosage compensation. In aphids, segregation distortion is already maximal (all sperm cells that do not carry an X degenerate, and those that are functional carry the single identical X chromosome), so no X-linked allele can further increase its transmission during spermatogenesis. Hence, there is no possibility for X-linked distorters to evolve, such that mechanisms to protect from distorters (i.e., sex-linked gene silencing) may have been lost.

In conclusion, we document an atypical genome-wide pattern of gene expression in aphids, with a high degree of masculinization of the X chromosome in both somatic and gonadic tissues. Our study reinforces the hypothesis that this masculinization evolved in response to sexual conflicts raised by the accumulation of male-beneficial alleles on the X (Jaquiéry et al. 2013). To further support this hypothesis, masculinization of the X should be assessed in distantly related aphid lineages (which diverged up to 200 MYA, Davis 2012) and other species that show an “aphid-like” life cycle and X-inheritance (e.g. *Strongyloides* nematodes, Nemetschke et al. 2010; Streit 2017). Finally, understanding the functional epigenetic or post-transcriptional mechanisms responsible for sex-biased gene expression in aphids would help to understand how such a strong chromosomal specialization in gene expression has been achieved.

## Methods

### Sex- and tissue-biased gene expression analysis

Gene expression levels in three different tissues (head, legs, gonads) of three reproductive morphs (male, sexual female, parthenogenetic female) were measured from RNA-Seq collected on a single pea aphid genotype (clone LSR1, from alfalfa, IAGC 2010). Aphids were reared on broad bean Vicia faba at less than five individuals per plant to prevent the production of winged morphs. Parthenogenesis was maintained under a 16-hour light regime and a temperature of 18°C. The production of sexual individuals was initiated by transferring larvae (at stage 3) to a 12-hour light regime at the same temperature of 18°C. Two generations later, sexual females and males were observed. A total of 100 adult parthenogenetic females (produced under 16-hour light regime), 100 adult sexual females and 100 adult males were immediately frozen into liquid nitrogen. Heads and legs were scalpel-cut. Twenty additional individuals per morph were also dissected in a saline solution with fine forceps to collect gonads. Gonads included testes and accessory glands in males and ovarioles in sexual females. Embryos of stage >10 (according to Miura et al. 2003) were removed from parthenogenetic female ovarioles (which already contain asexually developing embryos) to avoid the contribution of developing and late embryos to RNA production. All collected tissues were stored in RNA later (Qiagen) immediately after collection and pooled in batches before RNA extraction (with two replicates by sex and tissue). Hence, a total of 18 RNA extractions (3 morphs × 3 tissues × 2 biological replicates) were performed using the SV Total RNA Isolation System (Promega) according to manufacturer’s instructions. RNA quality was checked on Bioanalyzer (Agilent) and quantified on Nanodrop (Thermo Scientific). The 18 RNA samples were subsequently sent to the GetPlage platform (Toulouse, France) for library preparation (TruSeq Stranded mRNA Library Preparation kit) and 150 bp RNA paired-end sequencing (Illumina HiSeq3000).

After filtering for rRNA, reads from each library were mapped to the V2 assembly of the pea aphid genome (Acyr 2.0, Genbank accession GCA_000142985.2) using STAR version 2.5.2a (Dobin et al. 2013) with default parameters, except: outFilterMultimapNmax = 5, outFilterMismatchNmax = 3, alignIntronMin = 10, alignIntronMax = 50000 and alignMatesGapMax = 50000. Then, we counted reads mapped on exons of each predicted gene (NCBI Annotation release ID: 102) using FeatureCounts version 1.5.0-p3 (Liao et al. 2014) with default parameters except: -g gene -C -p -M --fraction. The numbers of mapped reads per library ranged from 19.8 to 28.6 million (mean 24.3 million, supplementary table S1).

We used the R package DESeq (Anders and Huber 2010) to normalize the libraries (upperquartile method with p = 0.75) and calculate CPM. Only genes with CPM>1 in at least two libraries (out of 18) were considered as expressed and retained in the analyses, unless mentioned otherwise. To identify genes predominantly expressed in a specific tissue or a morph, we imposed that a minimal percentage (70%) of the reads mapping to a given gene was sequenced from that specific tissue/morph. Doing so avoided the reliance on p-values from differential expression analyses (which in turn depend on the absolute expression level of a gene). This 70% threshold allowed identifying genes showing considerable bias in expression toward a tissue/morph, and to retrieve a large number of genes for more powerful analyses. So, when >70% of the reads mapping to a gene were from a given tissue from a given morph, the gene was classified as predominantly expressed in that tissue (e.g., PL+ for those predominantly expressed in legs of parthenogenetic females; etc). When >70% of the reads were from a given morph (but not restricted to a single tissue), the gene was classified as morph-specific (M+, F+ or P+, for males, sexual females, or parthenogenetic females, respectively). Similarly, genes mainly expressed (>70% of reads) in a tissue (but not restricted to a single morph) were classified as tissue-specific (G+, H+ or L+, for gonads, heads or legs, respectively). Expressed genes were thus classified into 16 mutually exclusive classes: MH+, ML+, MG+, FH+, FL+, FG+, PH+, PL+, PG+, M+, F+, P+, H+, L+, G+ or unbiased. Additional analyses with threshold values of 50%, 60%, 80% and 90% to identify a gene as predominantly expressed in a set of samples were also conducted to verify the robustness of our results to this parameter.

### Comparisons between chromosome types

The chromosomal location (X vs autosomes) of each gene was that of Jaquiéry et al. (2018). This represented the only assignments available at the start of the analyses. Since then, chromosome-level genome assemblies have been produced for the pea aphid (Li et al. 2019; Mathers et al. 2021). We found that our assignments to chromosomes were consistent for 97% to 98% of the genes located on the four mega scaffolds corresponding to the four pea aphid chromosomes, depending on the assembly. We therefore considered our initial assignment reliable. Furthermore, the number of NCBI predicted genes that we assigned to chromosome (18719) was larger than for the Li et al. (2019) assembly (17315). For the genome assembly from Mathers et al. (2021), DNA was obtained from a lineage sampled on Lathyrus, a host plant genus that harbors a cryptic species of the pea aphid complex (Peccoud et al. 2009; 2014) that is quite divergent from the LSR1 clone we used (Peccoud et al. 2009). Significant deviation from a random chromosomal distribution for each class of genes with a specific expression pattern was tested with two-sided binomial tests, the expected proportion was computed as the proportion of expressed genes on the X. To compare expression patterns between the three autosomes (supplementary table S4, supplementary figure S3), we used the assembly from Li et al. (2019). Expression breadth for X-linked and autosomal genes was estimated with τ (Yanai 2005) on log+1 transformed data.

### Investigation of gene families

To identify single-copy and multicopy genes in *A. pisum* genome, we used orthologs identified among 17 arthropod genomes by Boulain et al. (2018). Briefly, the longest protein isoform from each arthropod species was used to run OrthoDB_soft_1.6 (Kriventseva, et al. 2015) and the levels of orthology were assigned by referring to the species phylogeny established in Boulain et al. (2018). The groups of orthologs generated by OrthoDB were then used to identify *A. pisum* unique (single-copy) or duplicated (multicopy) genes. To control for possible differences in gene content between the X and autosomes, which could account for different expression patterns between chromosomes, we searched for gene families composed of two genes, one being on the X and the other on autosomes. To statistically compare frequencies of the various classes of expression between the X and autosomal copies with sufficient power given the limited sample size (n = 210 pairs of genes), the gene classes with male-biased expression (i.e., M+, MG+, ML+ and MH+ genes) were grouped into a single category of male-biased genes. We proceeded similarly for sexual female-biased genes and parthenogenetic female-biased genes. We also created a new class encompassing all genes that were not morph-biased (i.e, G+, H+, L+ and unbiased genes). For each aggregated gene class, we compared its frequency among chromosomes for single and duplicated copies and then among single and duplicated copies within chromosomes (chi-squared tests).

### Dosage compensation

To investigate dosage compensation, RPKM (reads per kilobase per million mapped reads) were calculated with EdgeR (Robinson et al. 2010), and only genes with RPKM > 1 in at least two of the 18 libraries were kept. After log-transformation of RPKM, differences in expression levels between chromosomes were examined with two-sided Mann-Whitney tests for each sex and each tissue. Then, the logarithm of the male to female (sexual or parthenogenetic) ratio of RPKM were estimated for X and autosomes for each tissue. As the uneven frequency of biased genes between chromosomes could interfere with dosage compensation patterns, the same analyses were performed by eliminating genes that showed a fold change in expression greater than 2 between males and females in at least one tissue.

## Acknowledgements

We are grateful to the Get-PlaGe platform of GenoToul (Toulouse, France) for transcriptome sequencing. We thank the recommenders Tanja Schwander and Charles Baer, as well as Ann Kathrin Huylmans and another anonymous referee for their constructive comments on previous drafts of this manuscript.

Preprint version 4 of this article has been peer-reviewed and recommended by Peer Community In Evolutionary Biology (https://doi.org/10.24072/pci.evolbiol.100148).

## Data, scripts and codes availability

The raw data are publicly available on NCBI (projects PRJNA547535 and PRJNA748848). The bioinformatic and R scripts, as well as the R input files can be found on Zenodo: https://doi.org/10.5281/zenodo.6242803.

## Supplementary material

**Supplementary table S1:**
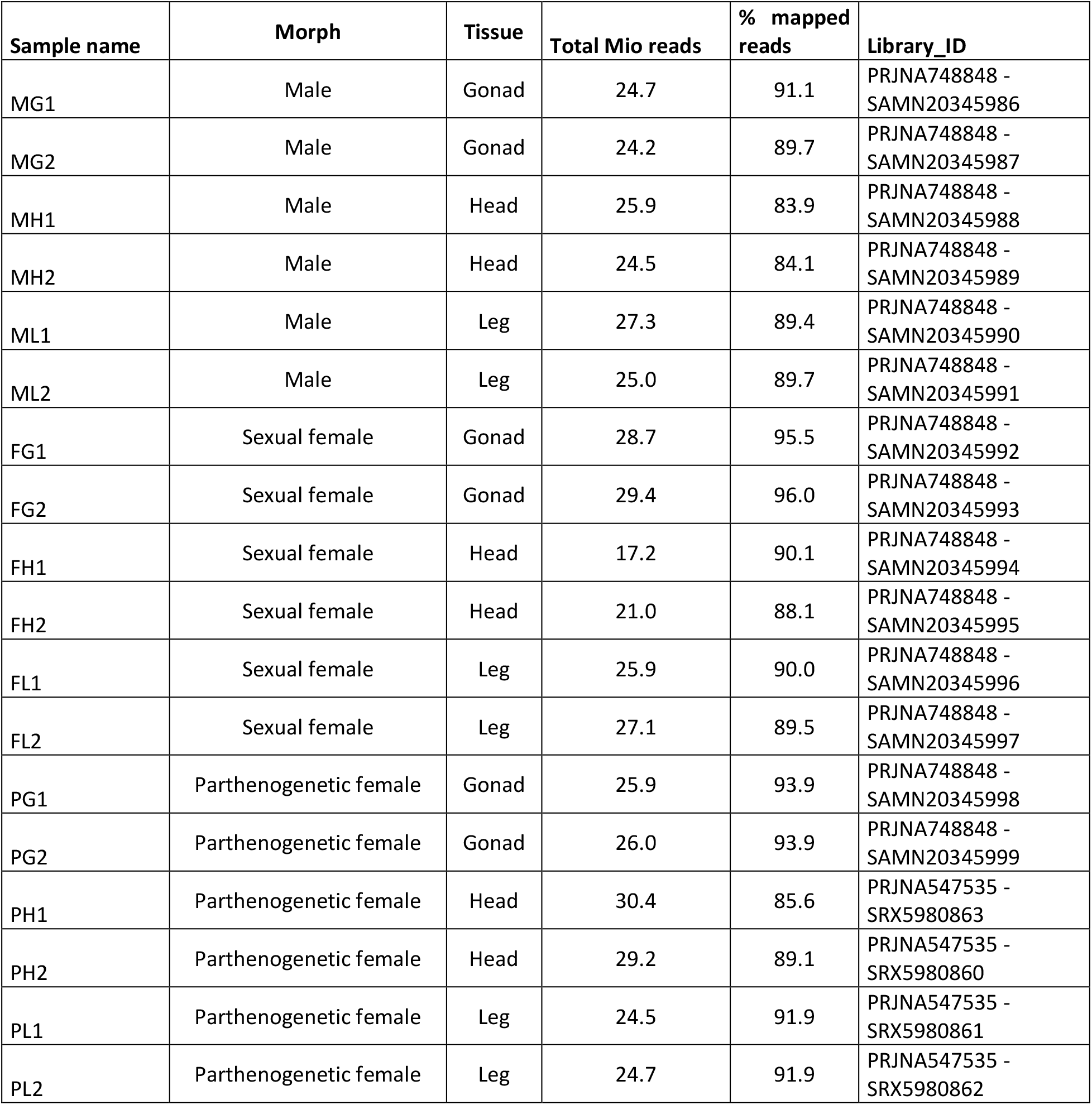
The different RNA sequencing libraries used in the present study.

| Sample name | Morph | Tissue | Total Mio reads | % mapped reads | Library_ID |
| --- | --- | --- | --- | --- | --- |
| MG1 | Male | Gonad | 24.7 | 91.1 | PRJNA748848 - SAMN20345986 |
| MG2 | Male | Gonad | 24.2 | 89.7 | PRJNA748848 - SAMN20345987 |
| MH1 | Male | Head | 25.9 | 83.9 | PRJNA748848 - SAMN20345988 |
| MH2 | Male | Head | 24.5 | 84.1 | PRJNA748848 - SAMN20345989 |
| ML1 | Male | Leg | 27.3 | 89.4 | PRJNA748848 - SAMN20345990 |
| ML2 | Male | Leg | 25.0 | 89.7 | PRJNA748848 - SAMN20345991 |
| FG1 | Sexual female | Gonad | 28.7 | 95.5 | PRJNA748848 - SAMN20345992 |
| FG2 | Sexual female | Gonad | 29.4 | 96.0 | PRJNA748848 - SAMN20345993 |
| FH1 | Sexual female | Head | 17.2 | 90.1 | PRJNA748848 - SAMN20345994 |
| FH2 | Sexual female | Head | 21.0 | 88.1 | PRJNA748848 - SAMN20345995 |
| FL1 | Sexual female | Leg | 25.9 | 90.0 | PRJNA748848 - SAMN20345996 |
| FL2 | Sexual female | Leg | 27.1 | 89.5 | PRJNA748848 - SAMN20345997 |
| PG1 | Parthenogenetic female | Gonad | 25.9 | 93.9 | PRJNA748848 - SAMN20345998 |
| PG2 | Parthenogenetic female | Gonad | 26.0 | 93.9 | PRJNA748848 - SAMN20345999 |
| PH1 | Parthenogenetic female | Head | 30.4 | 85.6 | PRJNA547535 - SRX5980863 |
| PH2 | Parthenogenetic female | Head | 29.2 | 89.1 | PRJNA547535 - SRX5980860 |
| PL1 | Parthenogenetic female | Leg | 24.5 | 91.9 | PRJNA547535 - SRX5980861 |
| PL2 | Parthenogenetic female | Leg | 24.7 | 91.9 | PRJNA547535 - SRX5980862 |

**Supplementary table S2:** Chromosomal location (X *vs* autosome) of the genes that show different patterns of expression.

| Pattern of expression | Abbreviation | Number of genes on the X chromosome | Number of genes on autosomes | Percentage of X-linked genes in each category | Percentage of all expressed X-linked genes | Percentage of all expressed autosomal genes | fold-change (X/A) |
| --- | --- | --- | --- | --- | --- | --- | --- |
| Unbiased genes | - | 1320 | 7526 | 15 | 43.4 | 69.1 | 0.6 |
| Head specific genes | H+ | 176 | 993 | 15 | 5.8 | 9.1 | 0.6 |
| Leg specific genes | L+ | 98 | 509 | 16 | 3.2 | 4.7 | 0.7 |
| Gonad specific genes | G+ | 113 | 398 | 22 | 3.7 | 3.7 | 1.0 |
| Male head specific genes | MH+ | 148 | 55 | 73 | 4.9 | 0.51 | 9.6 |
| Male leg specific genes | ML+ | 20 | 12 | 63 | 0.66 | 0.11 | 6.0 |
| Male gonad specific genes | MG+ | 637 | 533 | 54 | 20.9 | 4.9 | 4.3 |
| Male specific genes | M+ | 355 | 241 | 60 | 11.7 | 2.2 | 5.3 |
| Sexual female head specific genes | FH+ | 3 | 20 | 13 | 0.10 | 0.18 | 0.5 |
| Sexual female leg specific genes | FL+ | 11 | 34 | 24 | 0.36 | 0.31 | 1.2 |
| Sexual female gonad specific genes | FG+ | 119 | 256 | 32 | 3.9 | 2.4 | 1.7 |
| Sexual female specific genes | F+ | 8 | 50 | 14 | 0.26 | 0.46 | 0.6 |
| Parthenogenetic female head specific genes | PH+ | 8 | 66 | 11 | 0.26 | 0.61 | 0.4 |
| Parthenogenetic female leg specific genes | PL+ | 0 | 2 | 0 | 0.00 | 0.02 | 0.0 |
| Parthenogenetic female gonad specific genes | PG+ | 11 | 82 | 12 | 0.36 | 0.75 | 0.5 |
| Parthenogenetic female specific genes | P+ | 17 | 113 | 15 | 0.56 | 1.04 | 0.54 |
| Total (all expressed genes) | - | 3044 | 10890 | 0.22 | 100 | 100 | na |

The fold-change (rightmost column) is the ratio of the 6<sup>th</sup> column over the 7<sup>th</sup> column.

**Supplementary table S3:** Number of genes showing a given pattern of gene expression and their relative frequency, according to their chromosomal location and number of copies (for each two-copy-gene, one copy is on the X and the other on an autosome).

|  | X, single copy |  |  |  | A, single copy |  |  |  | X, two copies |  |  |  | A, two copies |  |  |  |
| --- | --- | --- | --- | --- | --- | --- | --- | --- | --- | --- | --- | --- | --- | --- | --- | --- |
| Pattern of expression* | # of genes |  | relative frequency |  | # of genes |  | relative frequency |  | # of genes |  | relative frequency |  | # of genes |  | relative frequency |  |
| M+ | 151 | 557 | 0.088 | 0.325 | 111 | 502 | 0.014 | 0.065 | 25 | 85 | 0.119 | 0.405 | 1 | 14 | 0.005 | 0.067 |
| MG+ | 341 |  | 0.199 |  | 364 |  | 0.047 |  | 49 |  | 0.233 |  | 13 |  | 0.062 |  |
| MH+ | 57 |  | 0.033 |  | 20 |  | 0.003 |  | 11 |  | 0.052 |  | 0 |  | 0.000 |  |
| ML+ | 8 |  | 0.005 |  | 7 |  | 0.001 |  | 0 |  | 0.000 |  | 0 |  | 0.000 |  |
| F+ | 4 | 65 | 0.002 | 0.038 | 31 | 189 | 0.004 | 0.025 | 0 | 13 | 0.000 | 0.062 | 0 | 5 | 0.000 | 0.024 |
| FG+ | 53 |  | 0.031 |  | 137 |  | 0.018 |  | 12 |  | 0.057 |  | 2 |  | 0.010 |  |
| FH+ | 1 |  | 0.001 |  | 9 |  | 0.001 |  | 0 |  | 0.000 |  | 0 |  | 0.000 |  |
| FL+ | 7 |  | 0.004 |  | 12 |  | 0.002 |  | 1 |  | 0.005 |  | 3 |  | 0.014 |  |
| P+ | 7 | 20 | 0.004 | 0.012 | 69 | 153 | 0.009 | 0.020 | 2 | 3 | 0.010 | 0.014 | 0 | 0 | 0.000 | 0 |
| PG+ | 9 |  | 0.005 |  | 61 |  | 0.008 |  | 0 |  | 0.000 |  | 0 |  | 0.000 |  |
| PH+ | 4 |  | 0.002 |  | 23 |  | 0.003 |  | 1 |  | 0.005 |  | 0 |  | 0.000 |  |
| PL+ | 0 |  | 0.000 |  | 0 |  | 0.000 |  | 0 |  | 0.000 |  | 0 |  | 0.000 |  |
| G+ | 48 | 1072 | 0.028 | 0.625 | 275 | 6824 | 0.036 | 0.889 | 17 | 109 | 0.081 | 0.519 | 15 | 191 | 0.071 | 0.91 |
| H+ | 102 |  | 0.060 |  | 648 |  | 0.085 |  | 1 |  | 0.005 |  | 4 |  | 0.019 |  |
| L+ | 65 |  | 0.038 |  | 316 |  | 0.041 |  | 4 |  | 0.019 |  | 2 |  | 0.010 |  |
| unbiased | 857 |  | 0.500 |  | 5585 |  | 0.728 |  | 87 |  | 0.414 |  | 170 |  | 0.810 |  |

\*See the methods section for a definition of the abbreviations. These different classes of genes were aggregated by morph (to sum all genes that show some bias in one of the three morphs, or that show no morph bias) and correspond to the data plotted on figure 3.

**Supplementary table S4:** Chromosomal location of genes that show different patterns of expression on the four chromosomes from the genome assembly of Li *et al.* (2019).

| Pattern of expression | Number of genes on: |  |  |  | Percentage of each category of gene on: |  |  |  | Fold-change between the X and each autosome: |  |  |
| --- | --- | --- | --- | --- | --- | --- | --- | --- | --- | --- | --- |
|  | X Scaffold_2 1773 | Autosome1 Scaffold_20 849 | Autosome 2 Scaffold_2 1967 | Autosome 3 Scaffold_2 1646 | X Scaffold_2 1773 | Autosome 1 Scaffold_2 0849 | Autosome 2 Scaffold_2 1967 | Autosome 3 Scaffold_2 1646 | X/Autosome 1 | X/Autosome 2 | X/Autosome 3 |
| **Unbiased genes** | 1248 | 3662 | 2525 | 1095 | 44.9 | 69.1 | 68.5 | 70.7 | 0.7 | 0.7 | 0.6 |
| **H+** | 156 | 498 | 344 | 114 | 5.62 | 9.39 | 9.33 | 7.35 | 0.6 | 0.6 | 0.8 |
| **L+** | 86 | 275 | 159 | 62 | 3.10 | 5.19 | 4.31 | 4.00 | 0.6 | 0.7 | 0.8 |
| **G+** | 108 | 194 | 137 | 53 | 3.89 | 3.66 | 3.71 | 3.42 | 1.1 | 1.0 | 1.1 |
| **MH+** | 131 | 28 | 25 | 5 | 4.72 | 0.53 | 0.68 | 0.32 | 8.9 | 7.0 | 14.6 |
| **ML+** | 14 | 4 | 2 | 5 | 0.50 | 0.08 | 0.05 | 0.32 | 6.7 | 9.3 | 1.6 |
| **MG+** | 558 | 274 | 188 | 72 | 20.10 | 5.17 | 5.10 | 4.65 | 3.9 | 3.9 | 4.3 |
| **M+** | 309 | 93 | 115 | 35 | 11.13 | 1.75 | 3.12 | 2.26 | 6.3 | 3.6 | 4.9 |
| **FH+** | 2 | 8 | 9 | 3 | 0.07 | 0.15 | 0.24 | 0.19 | 0.5 | 0.3 | 0.4 |
| **FL+** | 13 | 13 | 13 | 4 | 0.47 | 0.25 | 0.35 | 0.26 | 1.9 | 1.3 | 1.8 |
| **FG+** | 111 | 112 | 63 | 53 | 4.00 | 2.11 | 1.71 | 3.42 | 1.9 | 2.3 | 1.2 |
| **F+** | 7 | 24 | 14 | 6 | 0.25 | 0.45 | 0.38 | 0.39 | 0.6 | 0.7 | 0.7 |
| **PH+** | 8 | 26 | 30 | 8 | 0.29 | 0.49 | 0.81 | 0.52 | 0.6 | 0.4 | 0.6 |
| **PL+** | 0 | 2 | 0 | 0 | 0.00 | 0.04 | 0.00 | 0.00 | na | 0.0 | na |
| **PG+** | 9 | 35 | 29 | 13 | 0.32 | 0.66 | 0.79 | 0.84 | 0.5 | 0.4 | 0.4 |
| **P+** | 16 | 54 | 36 | 22 | 0.58 | 1.02 | 0.98 | 1.42 | 0.6 | 0.6 | 0.4 |
| **Total (all expressed genes)** | 2776 | 5302 | 3689 | 1550 | 100 | 100 | 100 | 100 | na | na | na |

**Supplementary figure S1:**
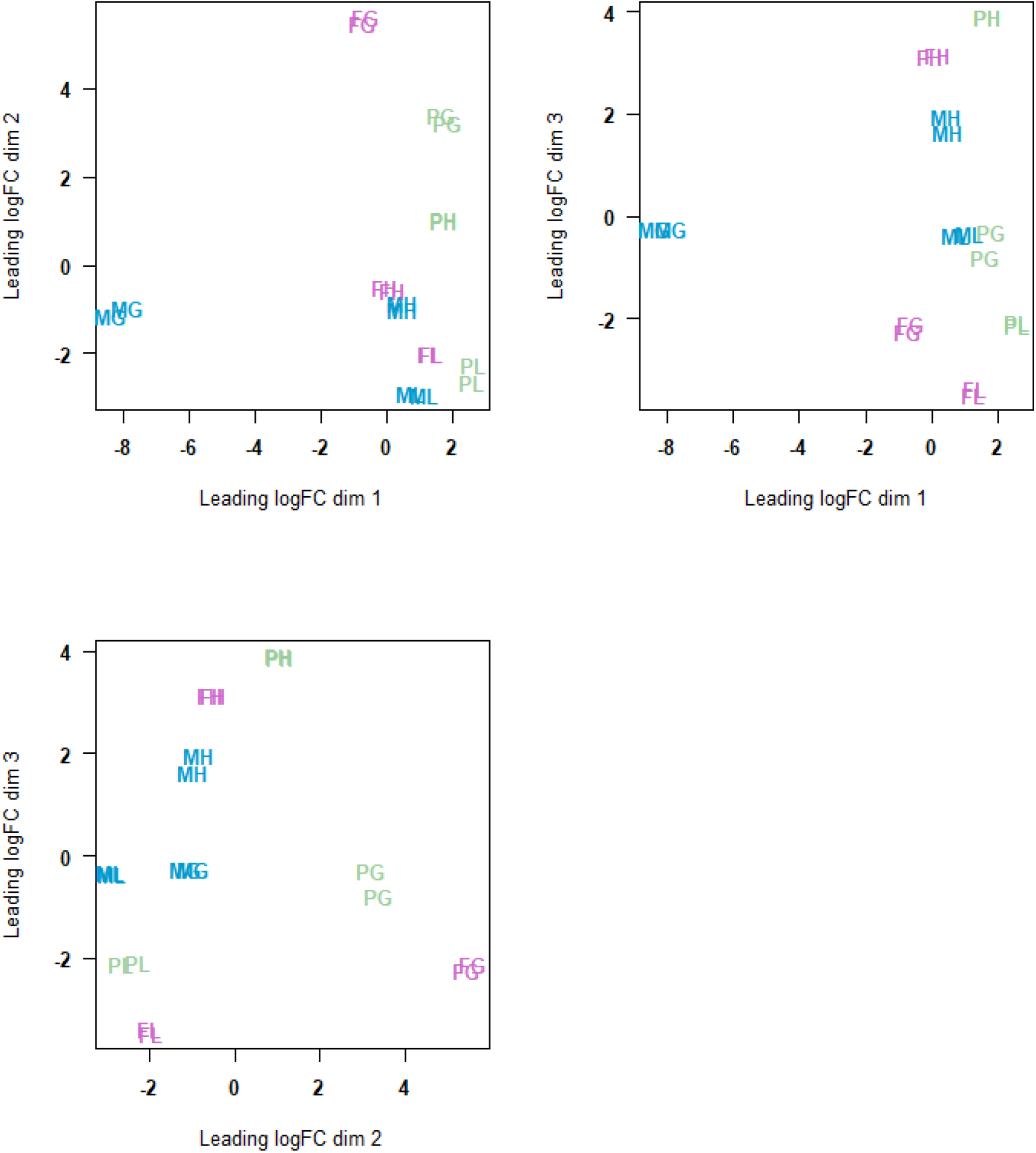
Multidimensional scaling (MDS, Limma R Package) of distances between gene expression profiles of the 18 samples using the plotMDS function with default settings (top = 500, gene.selection = “pairwise” in plotMDS function). MG: Male gonads, MH: male head, ML: male leg, FG: sexual female gonad, FH: sexual female head, FL: sexual female leg, PG: parthenogenetic female gonad, PH: parthenogenetic female head, PL: parthenogenetic female leg. The first dimension separates male gonads from all other samples, the second one separates mainly the female and parthenogenetic female gonads from legs and head samples, while the third one discriminates among head and leg samples.

**Supplementary figure S2:**
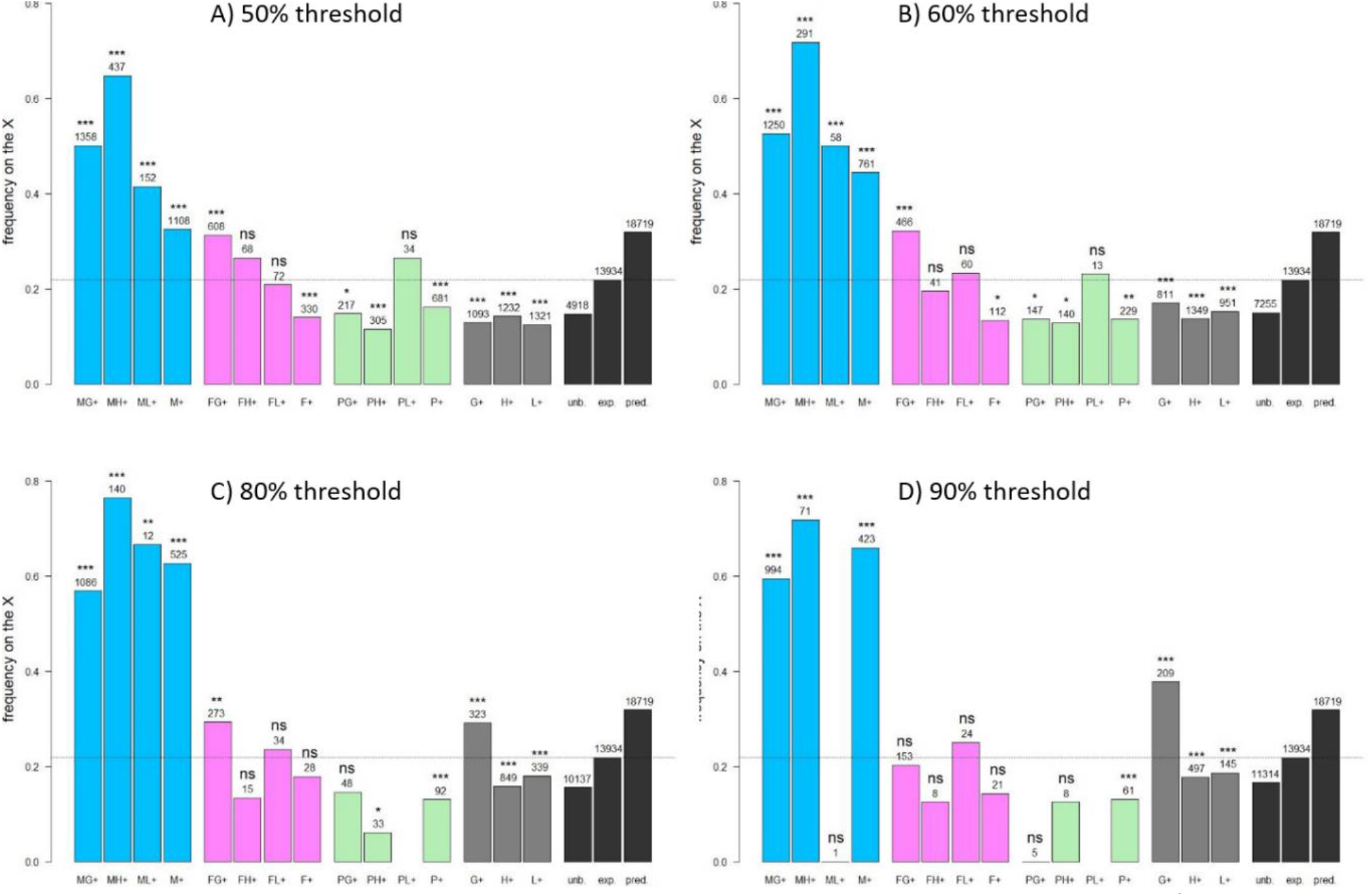
Proportions of X-linked genes among genes preferentially expressed in various morphs and/or tissues using different thresholds to define a gene as specific of a class of samples. Blue bars represent male-biased genes; MG+, MH+ and ML+: genes expressed preferentially in male gonads, heads and legs, respectively. M+: genes preferentially expressed in males when pooling all tissues, excluding the genes assigned to the previous categories. Pink bars represent sexual female-biased genes, with F standing for females and letters G, H and L having the same meaning as in males. Green bars represent parthenogenetic female-biased genes (P). Grey bars represent genes expressed preferentially in one of the tissues (gonads, heads or legs) and not limited to a particular morph. Black bars represent the frequency of X-linked genes among genes with unbiased expression (“unb”), genes expressed with CPM > 1 in at least two libraries (“exp”) or all predicted genes (“pred”). The horizontal dotted line represents the proportion of X-linked gene among expressed genes. The number of genes from each category is shown above bars, as well as the p-value (two-sided binomial tests against the expected frequency on the X chromosome estimated from expressed genes, which corresponds to the dotted horizontal black line). ***: *p* < 0.001; **: *p* < 0.01; *: *p* < 0.05; ns: *p* ≥ 0.05.

**Supplementary figure S3:**
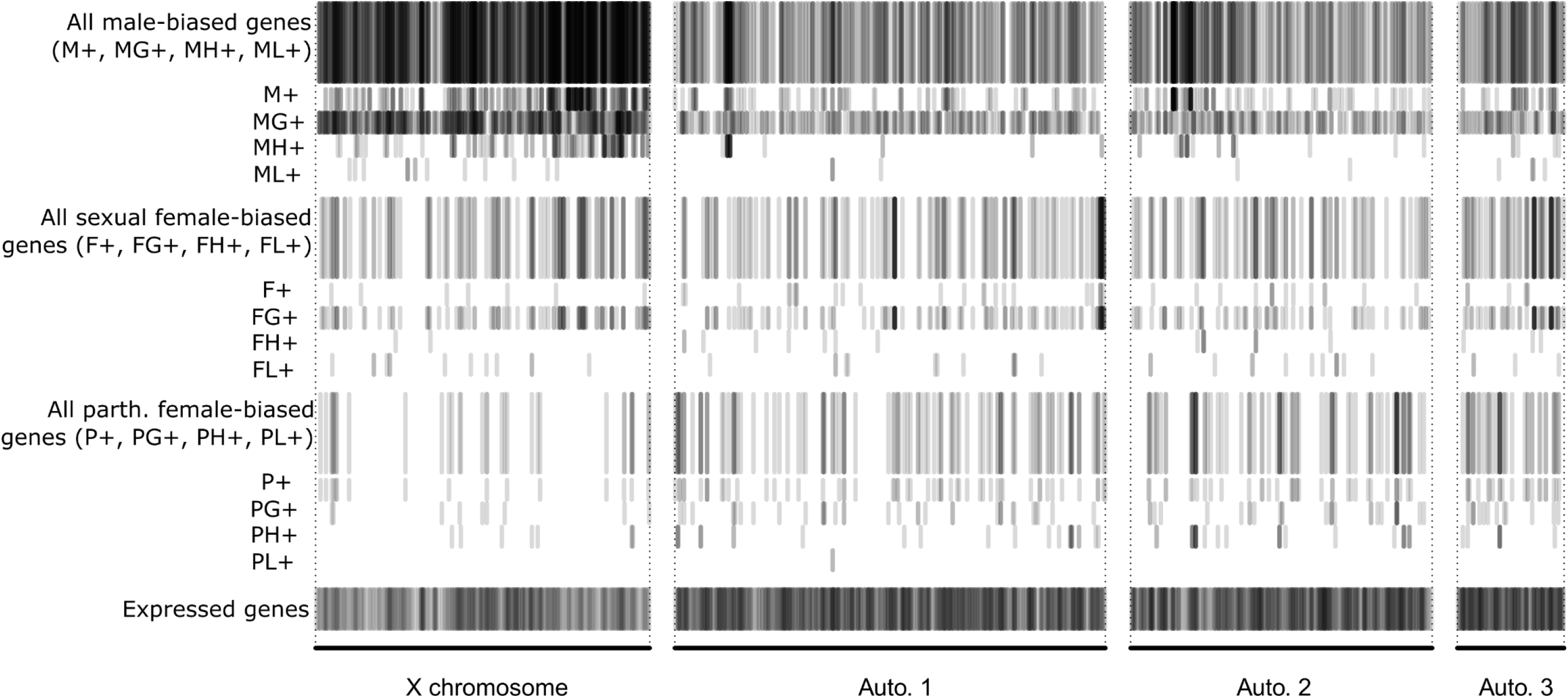
Distribution of the different types of sex-biased genes along the genome. Each gene with a biased expression is represented by a semi-transparent grayish bar, so that when many of such genes lay in the same genomic area, the region appears darker (e.g. the X chromosome harbors many male-biased genes). For each morph, the upper part of the graphic gathers all types of genes overexpressed in that morph (i.e. M+, MG+, MH+, ML+ for males) to get a global view. Just below, we then show the chromosomal position of these genes separately for each of the different classes of sex-biased genes. The chromosomal distribution of expressed genes is also presented (but with a lighter gray scale due to the large number of genes).

**Supplementary figure S4:**
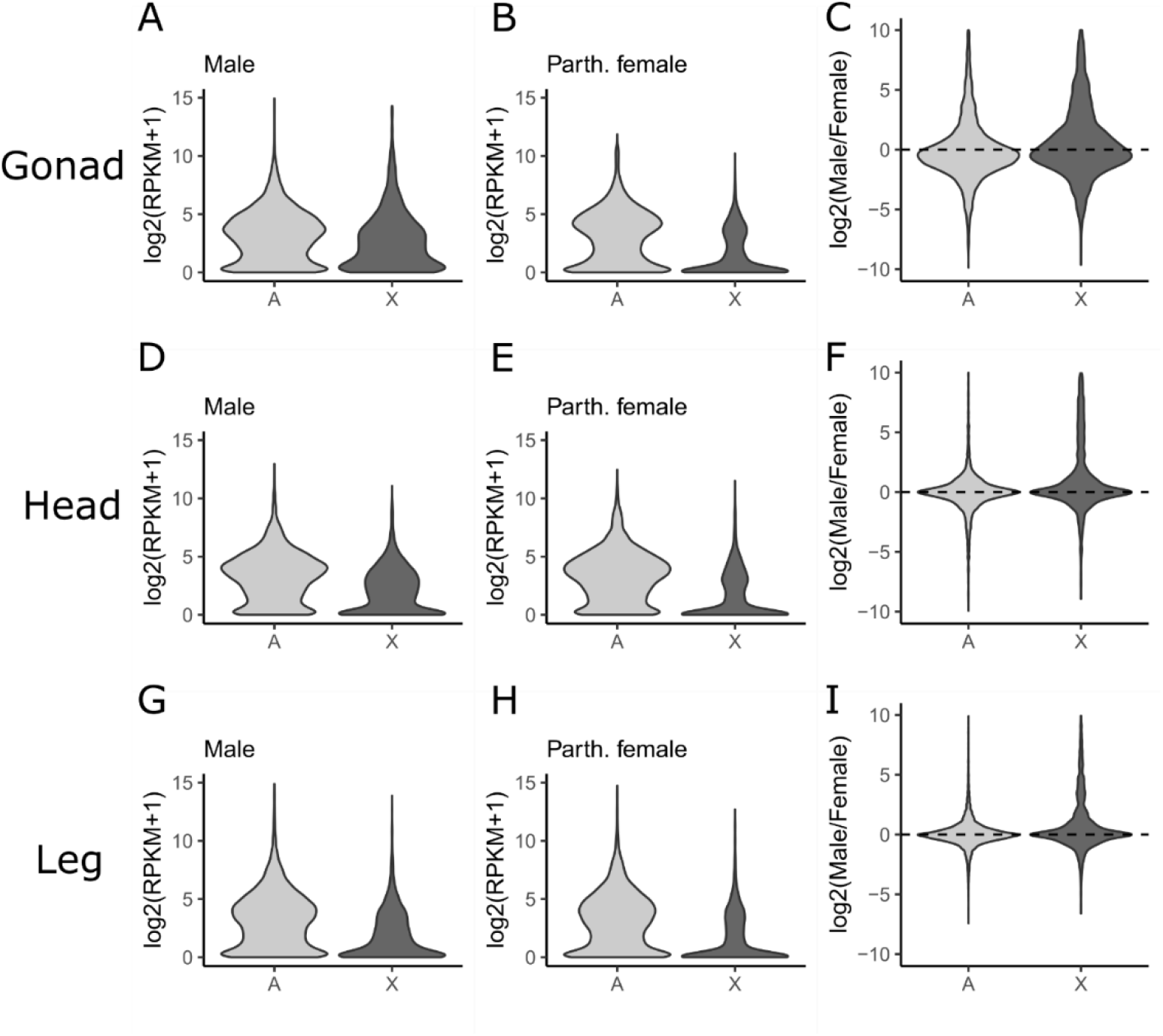
Logarithm of gene expression (RPKM+1) in males and parthenogenetic females for X-linked and autosomal genes in gonads (panels A and B), head (D, E) and legs (G, H). X-linked genes are significantly less expressed than autosomal genes in all cases (two-sided Mann-Whitney tests, *p* < 10^−15^). Logarithm of male to parthenogenetic female ratio of RPKM is also shown for the three tissues (panels C, F and I).

**Supplementary figure S5:**
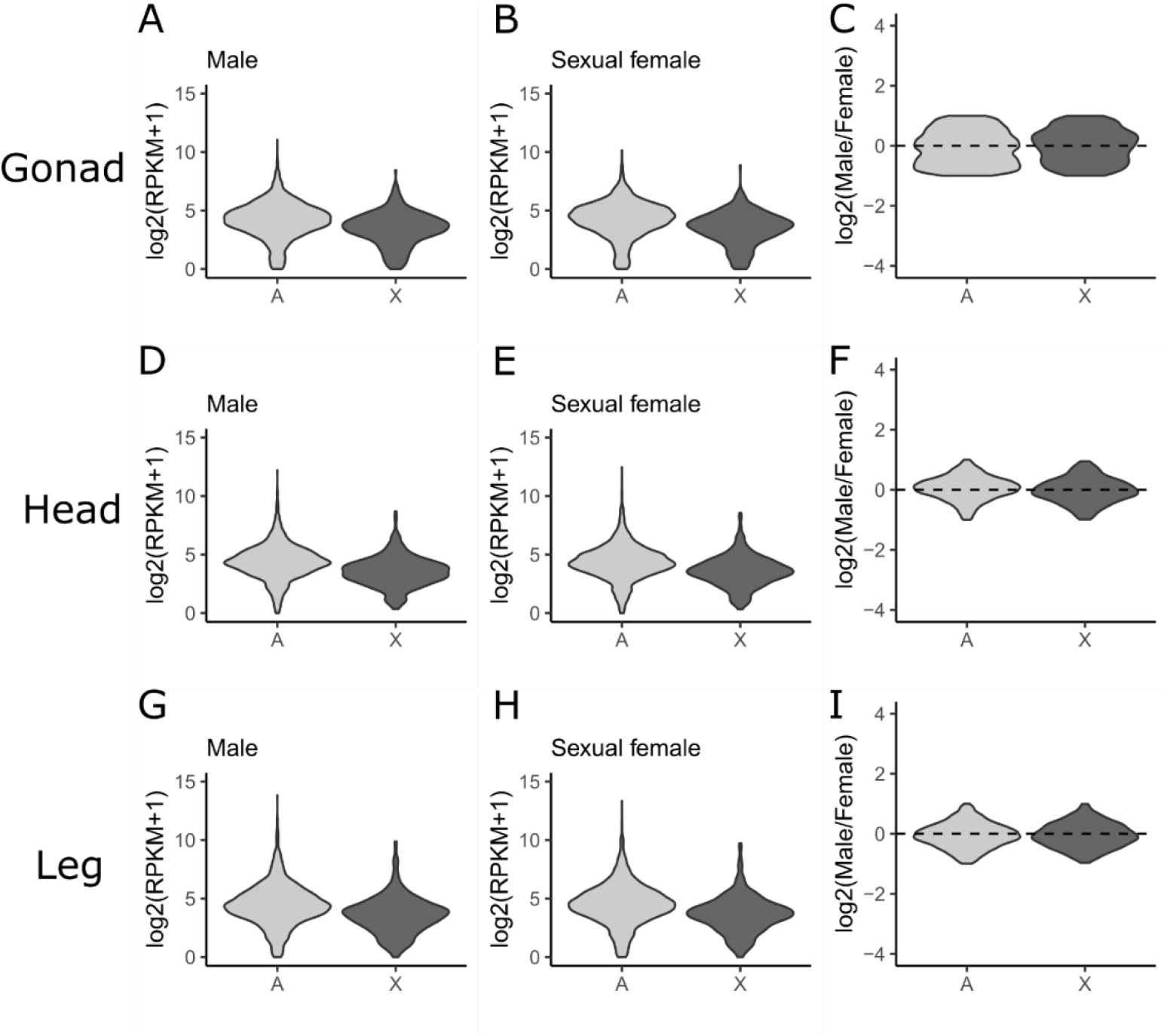
Logarithm of gene expression (RPKM+1) in males and sexual females for X-linked and autosomal genes in gonads (panels A and B), head (D, E) and legs (G, H). Only genes showing a less than 2-fold change in expression between males and sexual females within each of the tissues were retained. X-linked genes are significantly less expressed than autosomal genes in all cases (two-sided Mann-Whitney tests, *p* < 10^−15^). Logarithm of male to parthenogenetic female ratio of RPKM is also shown for the three tissues (panels C, F and I).

**Supplementary figure S6:**
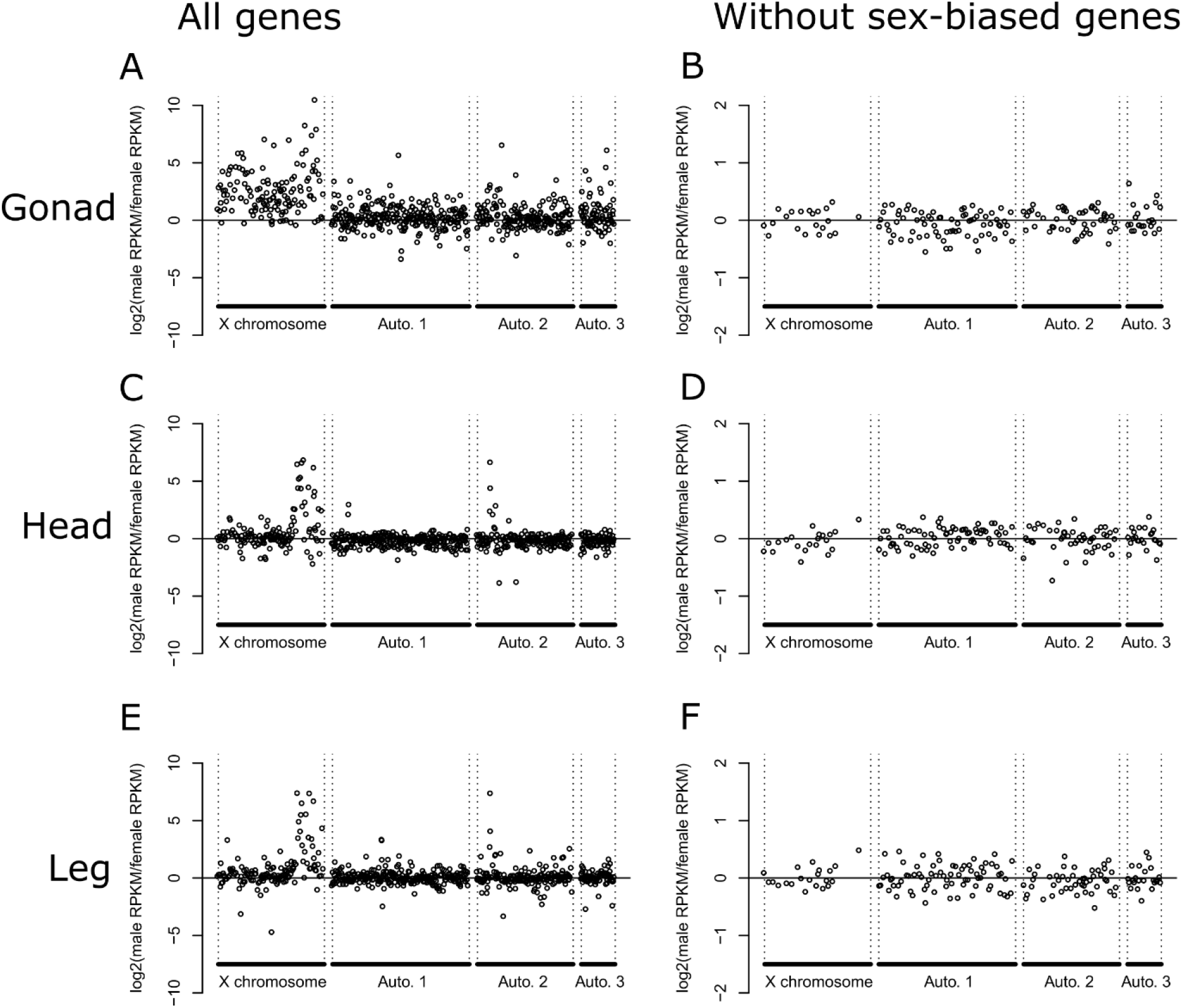
Logarithm of male to sexual female ratio of expression (in RPKM) along chromosomes in gonads (panels A and B), heads (C and D) and legs (E and F). In panels A, C and E, all expressed genes were included, while only genes with a less than 2-fold change in expression between males and sexual females within each of the tissues were retained in panels B, D and E (i.e. sex-biased genes were removed). To mitigate the high inter-gene variation, which could mask a pattern, we made sliding averages over 20 genes along the chromosomes (i.e. each point is the average over 20 genes).

**Supplementary text S1**: A large portion of genes were represented by less than 1 CPM in less than two (out of the 18 samples) and were classified as unexpressed in our 3-morph x 3-tissue RNAseq experiment. Unexpressed genes were particularly frequent on the X chromosome (see figure 1). To further investigate whether the higher proportion of genes classified as unexpressed on the X compared to autosomes emerges because a larger fraction of X-linked genes tend to be narrowly expressed (i.e. restricted to a particular unsampled tissue) or because a substantial fraction of X-linked genes are not expressed at all, we reanalyzed previous RNAseq data collected on whole body of males, sexual females and parthenogenetic females (3 morphs x 2 replicates, NCBI BioProject PRJNA209321, see Jaquiéry et al. 2013). The six RNAseq samples were re-analyzed as described in the main text for the tissue samples. Read counts per gene were then summed over the two replicates of the same morph (to have a per morph count), and also over the 6 samples (to get an overall expression count). We found that 40.3% of the X-linked genes considered as not expressed in the 3-morph x 3-tissue RNAseq experiment showed 0 reads over the 6 whole body samples, against 23% for the autosomal genes (see table A below).

**Table A:** Expression in whole-body samples of males, sexual females and parthenogenetic females for X-linked and autosomal genes that were considered as not expressed in the 3-morph x 3-tissue RNAseq experiment.

|  | Total number of reads over the 6 whole body samples |  |  |  |
| --- | --- | --- | --- | --- |
|  | 0 read | 1-5 reads | 6-20 reads | >20 reads |
| X linked - considered as not expressed (n=2917) | 40.3% | 34.4% | 18.9% | 6.3% |
| Autosomal - considered as not expressed (n=1868) | 23.0% | 26.6% | 28.9% | 21.5% |

We also found that the genes that we considered as expressed based on the 18 tissue samples were supported by high expression in whole body samples of the different morphs (see figure A below), while those considered as not expressed showed a much-reduced expression in all morphs.

**Figure A:**
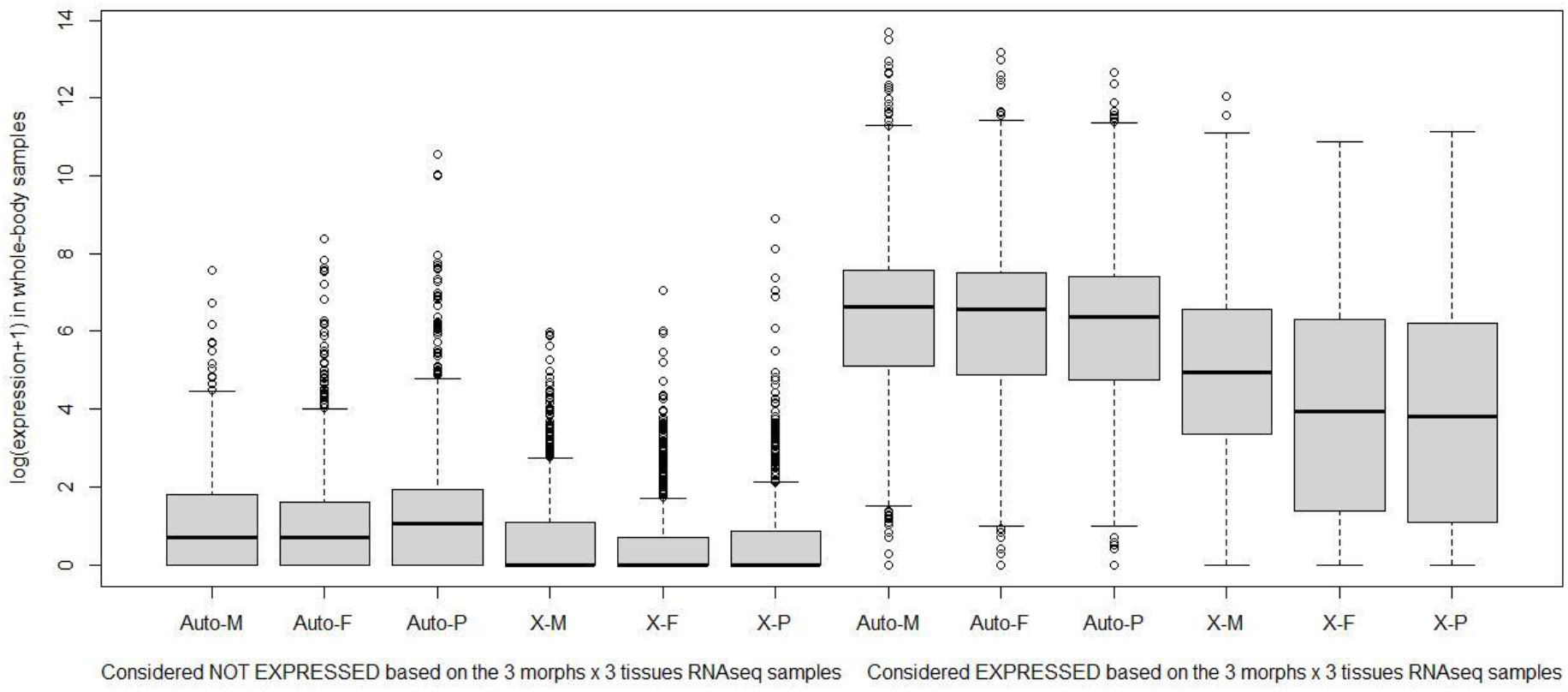
Gene expression in whole body samples of males (M), sexual females (F) and parthenogenetic females (P) as a function of the gene localization (X *vs* Autosomes) and of whether the gene is considered as expressed or not in the 3-morph x 3-tissue RNAseq experiment.

